# Electroporation-mediated delivery of protein biosensors for metabolic imaging in differentiated myotubes

**DOI:** 10.64898/2026.05.11.722572

**Authors:** Aki Kawamura, Cong Quang Vu, Nahoko Shimizu, Tsubasa Shibaguchi, Kazumi Masuda, Satoshi Arai

**Author notes:** Equal contribution (Aki Kawamura, Cong Quang Vu).

## Abstract

Understanding skeletal muscle metabolism involves real-time monitoring of key cellular parameters, such as calcium ions (Ca^2+^), adenosine triphosphate (ATP), cyclic adenosine monophosphate (cAMP), and intracellular temperature. Fluorescent protein (FP)–based biosensors are used for live-cell imaging of these signals with high spatiotemporal resolution. Differentiated myotubes are in vitro models used for physiological muscle metabolism research. However, efficient transfection of FP–based biosensors into these cells is challenging. Here, we developed an electroporation-based strategy for delivering recombinant protein biosensors into fully differentiated myotubes. Biosensors for Ca^2+^, ATP, cAMP, and temperature were recombinantly produced using *Escherichia coli* and introduced into myotubes using electroporation. Electroporation conditions were optimised to maximise delivery efficiency, preserve cell viability, and minimise cellular damage. We established a robust intracellular delivery system that effectively demonstrated Ca^2+^, ATP, and temperature dynamics. Furthermore, we achieved the successful co-delivery of two biosensors that enabled dual imaging of Ca^2+^ and cAMP in response to stimulation.

## Introduction

Real-time monitoring of cellular activities with high spatiotemporal resolution is essential for life science research. Fluorescent biosensors are used for real-time visualisation of signalling molecules and physicochemical parameters. Biosensors were initially small chemical indicators, with calcium probes, such as Fura-2 and Fluo-4.^1,2^ A major advantage of these chemical indicators is their experimental practicality; directly applying them to cells or tissues allows for immediate imaging examinations. However, the availability and applicability of these indicators, particularly for detecting signalling molecules and cellular metabolites, remain limited.^3,4^ This constraint reflects the difficulty of designing probes that achieve high selectivity for specific molecular targets. Fluorescent protein (FP)–based biosensors address this limitation by genetically encoding a fusion between a FP and target-specific binding domain, thereby achieving high specificity to the target molecule. Numerous biosensors capable of sensing various signalling molecules and metabolites were developed.^5^ Typically, FP–based biosensors are expressed in cells using gene delivery methods with transfection reagents, such as calcium phosphate, Lipofectamine, or polyethyleneimine.^6^ This approach works well for commonly used cell lines; however, certain cell types, including differentiated and primary cells, are often difficult to efficiently transfect. Viral infection methods can be used to overcome these limitations by using engineered viral vectors, such as adeno-associated virus or lentivirus, to deliver genes encoding biosensors into host cells or animals. However, these methods require specialised facilities, which can be inconvenient for researchers who are not routinely engaged in imaging studies.^4,7,8^ Both transfection and viral infection typically require 1–3 days to achieve sufficient protein expression, including the maturation time of FPs. To establish immediate effective imaging after delivery, purified FP–based biosensors would need to be directly applied to the biospecimen, similar to chemical indicators. This approach combines the molecular specificity of FP–based biosensors with the practicality of chemical indicators.

Protein delivery is a promising alternative to gene delivery. Unlike gene delivery, it is a rapid and simple approach used to directly introduce FP–based biosensors into cells. Moreover, it promotes uniform intracellular distribution of the introduced proteins, thereby alleviating issues associated with low expression efficiency and cell-to-cell variability. Several strategies have been developed for protein delivery, including the use of cell-penetrating peptide-, lipid-, and polymer-based protein transfection reagents; microinjection; and electroporation.^9^ In protein transfection, the reagents form complexes with the proteins that facilitate cellular uptake through direct membrane translocation or endocytosis. However, the delivered proteins are often trapped in endosomes,^10^ which limits their functional availability in the cytoplasm. Microinjection is used for precise delivery of defined amounts of proteins directly into the cytoplasm or nucleus of individual cells using a fine glass micropipette under a microscope. Although excellent spatial control and quantitative accuracy are achieved with microinjection, it is technically demanding, low throughput, and unsuitable for large-scale applications. In contrast, electroporation can be used to deliver proteins into cells by applying short electric pulses that transiently permeabilise the plasma membrane, allowing macromolecules to enter the cells. Therefore, high delivery efficiency is achieved in several cell types, including those that are difficult to transfect,^11^ making it preferrable for protein delivery.

In this study, we used electroporation to deliver purified proteins of FP–based biosensors into differentiated myotubes, which are difficult to transfect with conventional transfection reagents. We demonstrated the delivery of several biosensors for sensing calcium ions (Ca^2+^), adenosine triphosphate (ATP), cyclic adenosine monophosphate (cAMP), as well as intracellular temperature. We optimised electroporation conditions to maximise delivery efficiency, preserve cell viability, and minimise cellular damage. The functionality of the delivered biosensors was validated using fluorescence lifetime– and fluorescence intensity-based microscopy. Furthermore, we achieved dual imaging of Ca^2+^ and cAMP in differentiated myotubes, allowing the visualisation of these two elements within the same cells.

## Results

### Concept and design

We sought to establish a methodology for investigating muscle metabolism. Achieving this goal requires the ability to monitor key intracellular signals that coordinate energy production and contractile function. Among these, Ca^2+^ is a primary trigger of excitation-contraction coupling and regulate downstream metabolic pathways. ATP is the central energy currency that sustains muscle activity, and cAMP is an important second messenger controlling signalling pathways related to energy balance and cellular adaptation. Intracellular temperature is another critical parameter used to reflect metabolic activity and is essential to the processes, such as shivering and non-shivering thermogenesis.

Differentiated myotubes are physiological in vitro models used for muscle metabolism research. However, efficient delivery of biosensors into these cells remains challenging due to their low transfection efficiency. To establish efficient delivery of biosensors to myotubes, we developed an electroporation-based strategy for delivering recombinant protein biosensors: Tq-Ca-FLITS (Ca^2+^),^12^ MaLionG (ATP),^13^ Pink Flamindo (cAMP),^14^ and ELP-TEMP (temperature)^15^ into differentiated myotubes (**Fig. 1a**). Starting from the differentiation stage, fully differentiated myotubes were electroporated with recombinant biosensors on day 8, and intracellular protein stability was assessed on days 9 and 14 using microscopy (**Fig. 1b**). As a control, plasmids encoding the biosensors were electroporated into cells on day 3, allowing their expression during differentiation, and imaging was performed on days 9 and 14. Electroporation was performed using two approaches: (1) in adherent and (2) suspended myotubes using trypsin-mediated detachment. Hereafter, these conditions are referred to as “adherent” and “suspended”, respectively (**Fig. 1c**).

**Figure 1.**
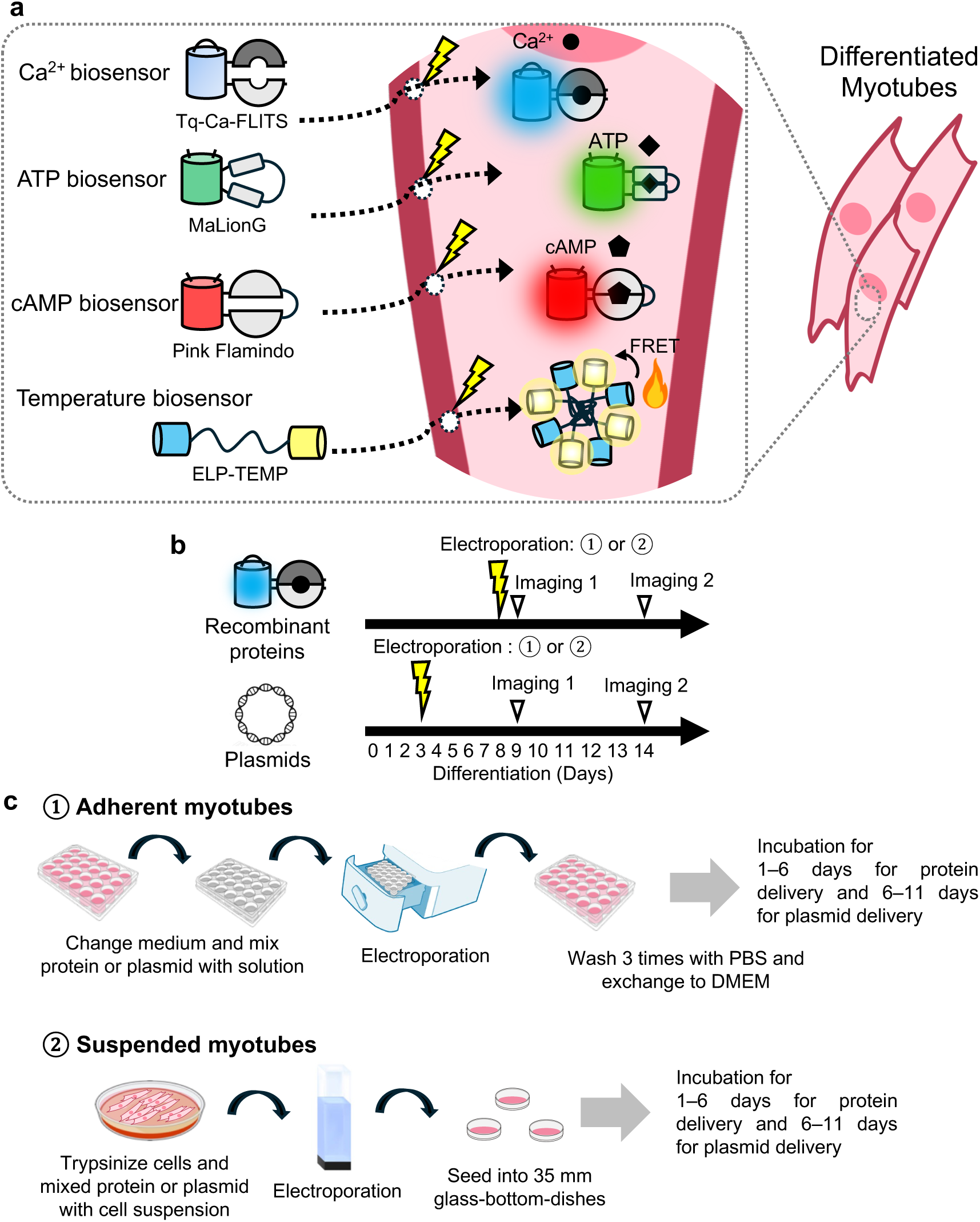
Concept and design of protein delivery of fluorescent protein–based biosensors into differentiated myotubes. (a) Schematic illustration of the experimental concept. Recombinant biosensors for Ca^2+^ (Tq-Ca-FLITS), ATP (MaLionG), cAMP (Pink Flamindo), and temperature (ELP-TEMP) are delivered into differentiated myotubes by electroporation, enabling real-time monitoring of multiple intracellular parameters, including Ca^2+^, ATP, cAMP, and temperature. (b) Experimental timeline for protein and plasmid delivery. Differentiated myotubes were electroporated with recombinant proteins on day 8, followed by imaging on days 9 and 14. For comparison, plasmids encoding the biosensors were introduced on day 3, allowing expression during differentiation, with imaging performed at the same time points. (c) Workflow of electroporation approaches. (1) Adherent myotubes: culture medium is replaced, biosensors (protein or plasmid) are added, followed by electroporation and washing. (2) Suspended myotubes: cells are detached by trypsinisation, mixed with biosensors, electroporated in suspension, and re-seeded onto glass-bottom dishes. Incubation times prior to imaging differ for protein (1–6 days) and plasmid (6–11 days) delivery. Illustration from NIH BIOART Source (https://bioart.niaid.nih.gov).

### Optimisation of electroporation conditions for protein and plasmid delivery in differentiated myotubes

We optimised the electroporation conditions using the Tq-Ca-FLITS biosensor by systematically varying the electric field strength during pulse stimulation. For adherent myotubes, electroporation was performed using programs CA-154, CA-215, FF-166, and FB-191, whereas for suspended myotubes, voltages ranging from 75 to 175 V were applied (**Fig. 2a**). After electroporation, fluorescence images were acquired on days 9 and 14 to evaluate delivery efficiency and signal stability **(Fig. 2b–e**).

**Figure 2.**
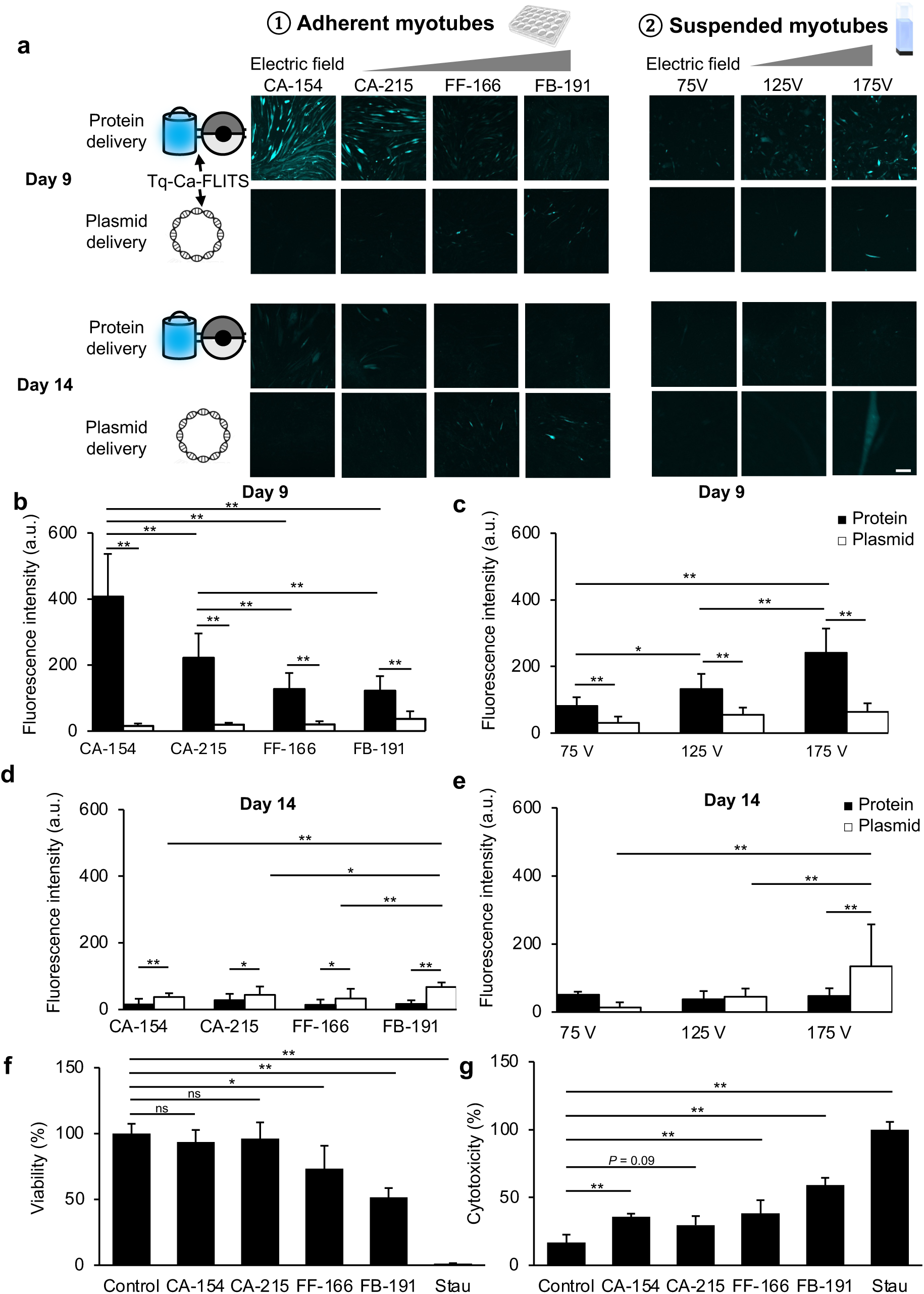
Optimisation of electroporation conditions for protein and plasmid delivery in differentiated myotubes. (a) Representative fluorescence images of Tq-Ca-FLITS delivered into differentiated myotubes under various electroporation conditions. Protein and plasmid delivery were compared in adherent myotubes using four programs (CA-154, CA-215, FF-166, and FB-191) and suspended myotubes at different voltages (75, 125, and 175 V). Images were acquired on day 9 and 14 after electroporation. Scale bar, 200 μm. (b, c) Quantification of fluorescence intensity on day 9 for adherent (b) and suspended (c) myotubes. Significantly higher fluorescence intensity was established with protein delivery than that with plasmid delivery across most conditions. (d, e) Quantification of fluorescence intensity at day 14 for adherent (d) and suspended (e) myotubes. Fluorescence intensity from protein delivery decreased over time and was lower than that from plasmid delivery at later time points. (f) Cell viability (%) of adherent myotubes following electroporation under different programs, compared with non-electroporated control and staurosporine (Stau)-treated cells (positive control). (g) Cytotoxicity (%) under the same conditions as in (f). Increased cytotoxicity occurred under strong electric fields. Data are presented as mean ± SD (*N* = 4). Significant interactions between electric fields (CA-154, CA-215, FF-166, and FB-191) and delivery methods (plasmid vs. protein), as well as significant main effects, were observed using two-way analysis of variance (ANOVA) followed by Bonferroni’s post hoc test for b–d. One-way ANOVA followed by Dunnett’s post-hoc test was performed for f and g. \**P* < 0.05; \*\**P* < 0.01; ns, not significant.

For adherent myotubes, significantly higher fluorescence intensity was achieved with protein delivery than that with plasmid delivery for all electroporation conditions on day 9 (CA-154: *P* < 0. 001, CA-215: *P* < 0. 001, FF-166: *P* < 0. 001, FB-191: *P* = 0.004) (**Fig. 2b**). Among the tested conditions, the highest fluorescence intensity occurred with the CA-154 program, the weakest electric field, indicating optimal delivery efficiency. Fluorescence intensity was sufficiently high on day 9 to enable robust imaging but substantially decreased by day 14. By day 14, fluorescence intensity in protein-delivered samples was significantly lower than that in plasmid-delivered samples for all conditions (CA-154: *P* = 0.007, CA-215: *P* = 0.043, FF-166: *P* = 0.026, FB-191: *P* < 0. 001) (**Fig. 2d**), indicating more rapid signal decay following protein delivery compared with that using plasmid-based expression. Consistently, the signal intensity increased with higher protein concentrations; significantly higher fluorescence was achieved with 300 μM than that with 3 μM (*P* < 0.001) and 30 μM (*P* < 0.001) under the CA-154 condition (**Supplementary Fig. S1**).

For suspended myotubes, a similar trend was observed. On day 9, significantly higher fluorescence intensity was achieved with protein delivery than that with plasmid delivery for all voltages (75 V: *P* = 0.003, 125 V: *P* < 0. 001, 175 V: *P* < 0. 001) (**Fig. 2c**), with the highest signal observed at 175 V. However, as in adherent myotubes, fluorescence from protein delivery substantially decreased by day 14 and remained significantly lower than that from plasmid delivery (175 V: *P* < 0. 001) (**Fig. 2e**). Overall, higher delivery efficiency was achieved with adhesion-based electroporation than that with suspension-based electroporation, which was mostly pronounced under weak electric fields. Notably, the enhanced efficiency observed at low electric fields in adherent myotubes was unexpected.

To evaluate potential cellular damage caused by electroporation, we assessed cell viability and cytotoxicity. In adherent myotubes, cell viability remained comparable to that of non-electroporated controls (100 ± 7.4%) under low electric fields, such as CA-154 (93.5 ± 9.1%) and CA-215 (96.2 ± 12.2%). However, cell viability significantly decreased under strong electric fields, such as FF-166 (73.3 ± 17.5%) (*P* = 0.020) and FB-191 (51.5 ± 7.1%) (*P* < 0.001) (**Fig. 2f**). Cytotoxicity was elevated under all electroporation conditions [CA-154: 35.9 ± 2.2%, *P* = 0.008; CA-215: 29.7 ± 6.6%, *P* = 0.090; FF-166: 38.2 ± 9.9%, *P* = 0.003 and FB-191: 59.1 ± 5.3%, *P* < 0.001] compared with that of the non-electroporated control (16.9 ± 5.7%). Notably, cytotoxicity exceeded 50% under the strongest electric field, FB-191 (*P* < 0.001) (**Fig. 2g**). Near-complete loss of viability (1.2 ± 0.3%) and maximal cytotoxicity (100.0 ± 5.7%) were observed in the 10-µM staurosporine positive control (*P* < 0.001) (**Fig. 2f,g**).

Our results indicate the highest fluorescence signal among all tested conditions was achieved with protein delivery at a concentration of 300 µM into adherent myotubes using the CA-154 electroporation condition. This method outperformed both suspension-based and plasmid delivery and maintained high cell viability. Therefore, it was used for all subsequent experiments.

### Applications of protein delivery for FP–based biosensors into differentiated myotubes

We delivered FP–based biosensors, including Tq-Ca-FLITS, MaLionG, ELP-TEMP, and Pink Flamindo, into differentiated myotubes to measure Ca^2+^, ATP, temperature, and cAMP, respectively. We investigated whether recombinant proteins produced in *Escherichia coli (E. coli)* retain their functionality within mammalian cells.

#### Ca^2+^ imaging with Tq-Ca-FLITS

Intracellular Ca^2+^ is pivotal to the initiation of muscle metabolism. The propagation of action potentials triggers the release of Ca^2+^ from the sarcoplasmic reticulum into the cytoplasm through the ryanodine receptor (RyR), a Ca^2+^ release channel.^16,17^ This process is initiated by stimuli, such as exercise, and the released Ca^2+^ plays a crucial role in maintaining muscle contraction. To monitor Ca^2+^ dynamics, we used Tq-Ca-FLITS, a fluorescence lifetime-based Ca^2+^ biosensor. Tq-Ca-FLITS consists of circularly permuted mTurquoise2 (mTQ2) fused to a Ca^2+^-sensing domain comprising of CaM and the M13 peptide. The fluorescence lifetime markedly increased when Tq-Ca-FLITS bound to Ca^2+^, as detected using quantitative fluorescence lifetime imaging microscopy (FLIM) imaging of intracellular Ca^2+^ dynamics (**Fig. 3a**). Following protein delivery with the optimised electroporation condition, clear fluorescence signals were observed using FLIM (**Fig. 3b–g**). Following the addition of 4 μM ionomycin, a Ca^2+^ ionophore, the fluorescence lifetime increased in the presence of extracellular Ca^2+^ in the imaging medium (**Fig. 3b,c**). The changes in the fluorescence lifetime were suppressed in the Ca^2+^-free medium, followed by a gradual decrease over time **(Fig. 3c**).

**Figure 3.**
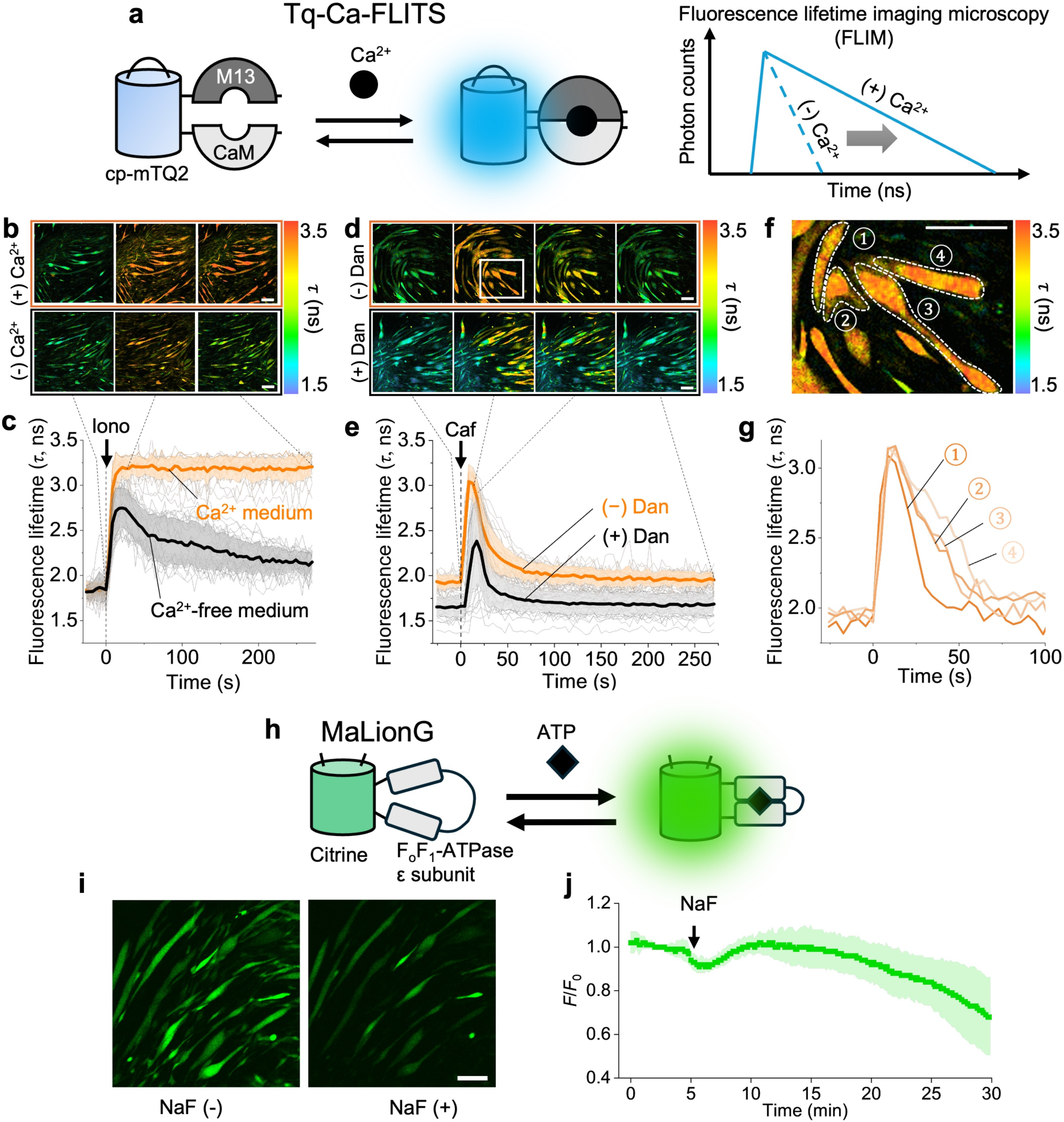
Calcium and ATP imaging of differentiated myotubes with Tq-Ca-FLITS, a fluorescence lifetime imaging microscopy (FLIM)–based Ca^2+^ biosensor and MaLionG, an intensity-based ATP biosensor. (a) Schematic illustration of the mechanism of Tq-Ca-FLITS. Tq-Ca-FLITS consists of circularly permuted mTurquoise2 (cp–mTQ2) fused with a Ca^2+^-sensing domain composed of calmodulin (CaM) and the M13 peptide. Upon binding to Ca^2+^, Tq-Ca-FLITS undergoes a conformational change, resulting in an increase in fluorescence lifetime, which is measured using FLIM). (b) FLIM images of differentiated myotubes delivered with Tq-Ca-FLITS proteins via electroporation and 4 μM ionomycin (Iono). (c) A graph showing fluorescence lifetime as a function of time under treatment with 4 μM Iono, with and without Ca^2+^ in the medium. (d) FLIM images of differentiated myotubes delivered with Tq-Ca-FLITS proteins via electroporation under treatment of 5 mM caffeine (Caf). (e) A graph showing fluorescence lifetime as a function of time under treatment with 5 mM Caf, with and without a 50 μM dantrolene (Dan) pretreatment. Data are show as mean ± SD (*n* = 12 cells, *N* = 3). (f,g) Heterogeneity in caffeine-induced responses among individual myotubes. (h) Schematic illustration of MaLionG. MaLionG consists of citrine fused with the F_0_F_1_-ATPase ε subunit, an ATP binding domain. Upon binding to ATP, MaLionG undergoes a conformational change, resulting in an increase in fluorescence intensity. (i) Fluorescence images of differentiated myotubes delivered with MaLionG protein via electroporation without and with NaF treatment. (j) Normalised fluorescence intensity (*F*/*F*_0_) of MaLionG with 100 mM NaF against time at 5 min. Data show mean ± SD (*n* = 12 cells, *N* = 3). Scale bars, 200 μm.

To induce RyR-mediated Ca^2+^ release and recapitulate muscle contraction signalling, we stimulated differentiated myotubes containing Tq-Ca-FLITS using caffeine.^18,19^ Overall, the fluorescence lifetime rapidly increased after adding 5 mM caffeine, then slowly returned to the basal levels (**Fig. 3d,e**). However, heterogenous responses were observed with the florescence lifetime in individual myotubes after caffeine stimulation (**Fig. 3f,g**). The observed Ca^2+^ dynamics are consistent with previous studies reporting Ca^2+^ changes during electrically evoked muscle contractions.^20,21^ The elevation of Ca^2+^ was minimally prevented with 50 μM dantrolene, an inhibitor of the RyR that suppresses Ca^2+^ release from the sarcoplasmic reticulum (**Fig. 3d,e**). Notably, the baseline fluorescence lifetime after treatment with dantrolene was lower than that before treatment, suggesting a reduction in cytoplasmic Ca^2+^ levels. This decrease likely reflects inhibition of Ca^2+^ release from the endoplasmic reticulum by inhibiting RyR1. The incomplete inhibition is consistent with previous studies reporting that RyR1 is not fully blocked by dantrolene but rather exhibits reduced open probability of the ion channel.^22^

#### ATP imaging with MaLionG

Monitoring ATP levels in muscle cells is essential, as ATP is the energy currency in cellular metabolism. To monitor ATP dynamics, we electroporated differentiated myotubes to deliver MaLionG, a fluorescence intensity-based ATP biosensor (**Fig. 3i**). MaLionG consists of citrine fused to the ε subunit of F_0_F_1_–ATPase, the ATP-binding domain. Upon ATP binding, the ε subunit undergoes a conformational change, leading to an increase in fluorescence intensity, characteristic of a conventional intensity-based indicator (**Fig. 3h**). We validated the working performance of MaLionG using 100 mM NaF, an enolase inhibitor. The normalised fluorescence intensity of MaLionG decreased within 30 min (**Fig. 3j**), consistent with ATP depletion due to inhibition of the glycolytic pathway.

#### Temperature imaging with ELP-TEMP, a Förster resonance energy transfer (FRET)–based temperature biosensor

The delivery procedure was optimised using Tq-Ca-FLITS, which contains a single FP in its construct. Therefore, we examined whether this approach is also applicable to FP–based biosensors with larger molecular sizes, particularly those comprising two FPs, such as Förster resonance energy transfer (FRET)–based biosensors. ELP-TEMP, a FRET–based temperature biosensor was used. The molecular weight of ELP-TEMP, including His-tag, is approximately 109 kDa, which is approximately two-fold larger than that of Tq-Ca-FLITS (∼51 kDa) and MaLionG (∼46 kDa). Monitoring intracellular temperature is essential for studies on shivering and non-shivering thermogenesis in muscle physiology. Additionally, in bioengineering, mild heating approaches have been used to induce muscle contraction and promote cellular biogenesis.^23^ Notably, optical heating using a near-infrared laser effectively manipulates muscle function at high spatiotemporal resolution.^24^

ELP-TEMP comprises a FRET pair of mTQ2 and mVenus (mV) fused to opposite ends of an ELP with temperature-dependent behaviour. Below the transition temperature (*T*_t_), the conformation of the ELP in the fusion protein is largely extended in solution, resulting in low FRET efficiency from mTQ2 to mV. In contrast, above the *T*_t_, the ELP moiety undergoes a conformational change, compacting into aggregates, reducing the distance between mTQ2 and mV, and thereby increasing FRET efficiency. Importantly, FRET–based biosensors can, in principle, be analysed using FLIM (i.e. FRET–FLIM). An increase in FRET efficiency upon temperature elevation led to a corresponding decrease in the fluorescence lifetime of the donor, mTQ2, as measured using FLIM (**Fig. 4a**). We evaluated whether ELP-TEMP functions in differentiated myotubes. Following electroporation, we observed the fluorescence from ELP-TEMP in the myotubes. To validate its temperature responsiveness, we applied controlled heating using confocal microscopy with a 1460-nm laser, which photothermally heated the surrounding medium (**Fig. 4b**). The fluorescence lifetime of mTQ2 decreased with increasing laser power (16.2, 25.6, 34.5, and 67.4 mW) (**Fig. 4c,d**), indicating that it could be used to detect temperature increments induced by photothermal heating.

**Figure 4.**
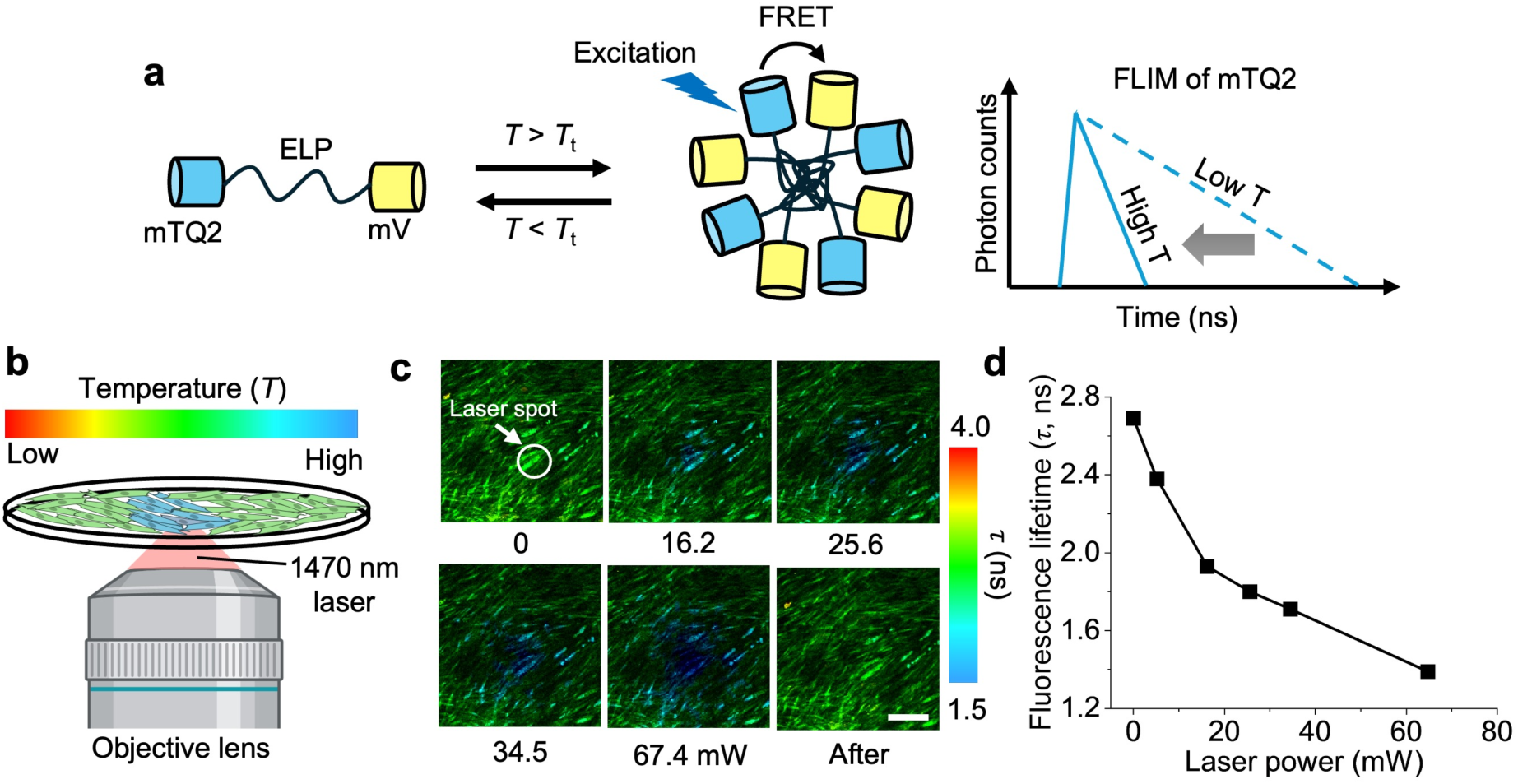
Temperature imaging of differentiated myotubes with ELP-TEMP, a Förster resonance energy transfer (FRET)–based temperature biosensors. (a) Schematic illustration of the temperature-sensing mechanism of ELP-TEMP. ELP-TEMP consists of mTurquoise2 (mTQ2) and mVenus (mV) fused to opposite ends of an ELP that is sensitive to temperature. Below the transition temperature (*T*_t_), the conformation of the ELP in the fusion protein is largely extended and dispersed in solution, resulting in low FRET efficiency from mTQ2 to mV. Above the *T*_t_, the ELP moiety undergoes a conformational change, compacting into aggregates, reducing the distance between mTQ2 and mV, and thereby increasing FRET efficiency. This increase in FRET efficiency leads to a decrease in the fluorescence lifetime of mTQ2, as measured using FLIM. (b) Schematic illustration of optical heating using 1470 nm laser. (c) Fluorescence lifetime images of ELP-TEMP at different laser power. (d) A graph of fluorescence lifetime of mTQ2 in ELP-TEMP at the laser spot after 1470-nm laser power. Scale bar, 200 μm. Illustrations were created with BioRender.com.

#### Dual imaging of Ca^2+^ and cAMP with Fast-GCaMP6f-RS09 and Pink Flamindo

We examined whether two different biosensors could be simultaneously delivered into differentiated myotubes. We performed fluorescence intensity-based dual imaging using Fast-GCaMP6f-RS09^25^ as a Ca^2+^ biosensor (green channel) and Pink Flamindo as a cAMP biosensor (red channel) because Ca^2+^ and cAMP signalling are closely associated in muscle physiology. For instance, Ca^2+^ influx activates Ca^2+^-dependent adenylyl cyclase, thereby promoting cAMP production. We focused on Ca^2+^, a skeletal muscle contraction mediator, and cAMP, a muscle physiology regulator, as two functionally complementary second messengers. Additionally, caffeine elevates cAMP levels and activates the cAMP response element–binding protein (CREB) signalling pathway in muscle cells.^26^

Fast-GCaMP6f-RS09 consists of a circularly permuted EGFP fused with calmodulin (CaM) and the M13 peptide (**Fig. 5a**), and Pink Flamindo is composed of mApple fused to mEpac1 (**Fig. 5b**). Fluorescence intensity is enhanced upon binding of the biosensors to the analytes, Ca^2+^ and cAMP, respectively. As the single FP–based biosensor was delivered at 300 µM, for dual imaging, Fast-GCaMP6f-RS09 and Pink Flamindo biosensors were delivered at 150 µM each, for a total concentration of 300 µM. After the co-delivery, clear fluorescence signals were observed from both biosensors (**Fig. 5c**). Upon stimulation with 5 mM caffeine, the fluorescence intensity of Fast-GCaMP6f-RS09 transiently increased, followed by a slight increase in the fluorescence of Pink Flamindo (**Fig. 5d**). Although the synchronous response with Ca^2+^ and cAMP remains unclear, the results demonstrate the feasibility of dual imaging using two FP–based biosensors.

**Figure 5.**
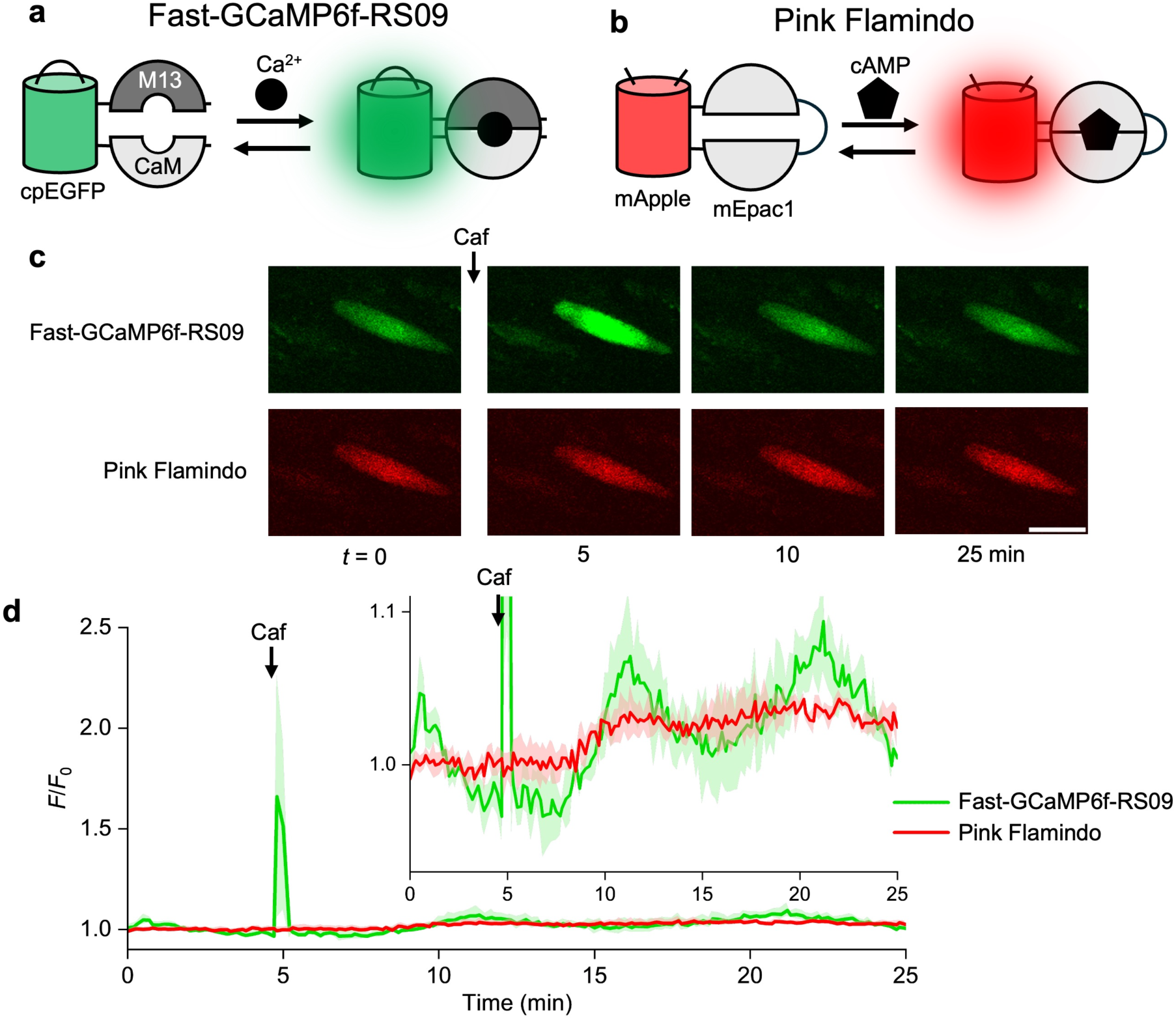
Ca^2+^ and cAMP dual imaging of differentiated myotubes with Fast-GCaMP6f-RS09 and Pink Flamindo. (a) Schematic illustration of the working mechanism of Fast-GCaMP6f-RS09. Fast-GCaMP6f-RS09 consists of circularly permuted enhanced green fluorescent protein (cpEGFP) fused with calmodulin (CaM) and the M13 peptide. Upon binding to Ca^2+^, Fast-GCaMP6f-RS09 undergoes a conformational change, resulting in an increase in fluorescence intensity. (b) Schematic illustration of the working mechanism of Pink Flamindo. Pink Flamindo consists of mApple fused with mEpac1, a cAMP binding domain. Upon binding to cAMP, Pink Flamindo undergoes a conformational change, resulting in an increase in fluorescence intensity. (c) Fluorescence images of Fast-GCaMP6f-RS09 and Pink Flamindo and (d) normalised fluorescence intensity (*F*/*F*_0_) of Fast-GCaMP6f-RS09 and Pink Flamindo with 5 mM caffeine (Caf) over time at 5 min. The inset shows a reduced scale. Data shown are mean ± SD (*n* = 4 cells). Scale bar, 200 μm.

## Discussion

This study demonstrates that FP–based biosensors can be efficiently delivered as recombinant proteins into adherent differentiated myotubes using electroporation. Electroporation has been used for various adherent cell types,^27,28^ and its potential for protein delivery^29^ and macromolecules^30^ has been suggested. However, its application for FP–based biosensor delivery in skeletal muscle cells has not been fully established. To address this gap, we delivered several FP–based biosensors for monitoring Ca^2+^, ATP, cAMP, and temperature via electroporation. We further demonstrated that these biosensors recombinantly produced in *E. coli* retained their functionality within mammalian cells and enabled real-time imaging of these intracellular signals (**Fig. 3–5**).

The optimised protocol in this study was used for efficient delivery into densely packed, mature myotubes after 9 days of differentiation. Notably, protein delivery was achieved under a relatively low electric field, which maintained cell viability and minimized cytotoxicity. To further evaluate the impact of electroporation on cell integrity, we compared weak (CA-154) and strong (FB-191) electric fields using western blotting. The total protein concentration and desmin, a cytoskeletal protein, remained comparable to non-electroporated controls, whereas both were markedly reduced under a strong electric field (FB-191) (**Supplementary Fig. S2a, b, d**). In contrast, no significant differences were observed in MyHC, a contractile protein, between electroporated and non-electroporated cells or weak and strong electric fields (**Supplementary Fig. S2c**). As mentioned earlier, the delivery efficiency was higher under weaker electric fields than that under strong electrical fields. Electroporation efficiency is determined by a balance between membrane permeabilisation and cell viability.^31^ In our study, sufficient and relatively homogeneous membrane permeabilisation across cells was achieved in differentiated myotubes under adherent conditions, confirming the high efficiency at relatively weak electric field strengths. This may preserve the cellular integrity, including efficient membrane resealing, which allows improved retention of the biosensor protein, consistent with **Supplementary Fig. S2c**. Although key cytoskeletal proteins were maintained under the CA-154 program, further studies with larger sample sizes will be required to evaluate the potential effects on other proteins, including those related to muscle function and mitochondrial activities.

Several limitations should be addressed. First, the method in this study involved the evaluation of cell viability, cytotoxicity, and protein integrity of mature myotubes (day 9); however, optimal electroporation conditions may vary depending on the maturation state of the cells. Therefore, protocols will need to be optimised for the specific characteristics of the target cell population. Second, the delivery method was initially optimised using a single FP–based biosensor Tq-Ca-FLITS and successfully extended to a FRET–based biosensor and co-delivery of two biosensors. However, the delivery method should be further optimised for each biosensor, particularly considering differences in molecular size and surface charge.

The proposed protein delivery method enables rapid imaging after the delivery. In contrast to plasmid delivery, which typically requires 1–3 days for the sufficient expression of FP–based biosensors, this approach can be conducted within 10–15 min after electroporation (**Supplementary Fig. S3**), thereby avoiding the loss of critical observation times for transient cellular states. Another key advantage is that cells remain attached to the dish during electroporation under adherent conditions, even after multiple washing steps. This allows for a large number of cells to be retained and analysed within a single field of view. Cell density is a critical factor in muscle research that requires the simultaneous observation of multiple myotubes is often required. Conversely, suspension-based protocols for myotube collection^32^ tend to result in a higher prevalence of myoblasts and immature myotubes rather than differentiated myotubes. Myoblasts and immature myotubes express fewer cytoskeletal proteins and receptors than myotubes,^33,34^ which limits their utility as models that accurately mimic skeletal muscle metabolism. Additionally, the adherent approach eliminates the need for reattachment and overnight incubation. Collectively, the combination of FP–based biosensor protein delivery and adherent myotube–based electroporation is a simple and convenient method for skeletal muscle physiology research.

We used several FP–based biosensors to compare two imaging modalities: conventional fluorescence intensity–based imaging and FLIM. Fluorescence lifetime is not related to factors, such as an expression level, focus drift, and excitation laser power, therefore, FLIM is used for a more quantitative readout, unlike conventional intensity-based imaging.^35^ Although FLIM has advantages for quantitative analysis, including FRET–FLIM, its broader application remains limited by the accessibility of instrumentation and relatively lower temporal resolution. Therefore, approaches based on conventional intensity-based imaging are preferred. In this study, we successfully achieved real-time monitoring of ATP depletion and captured the dynamics of Ca^2+^ and cAMP using intensity-based biosensors. Additionally, the Ca^2+^ level, which cannot be reliably extracted from an intensity-based analysis, was established using the fluorescence lifetime–based biosensor, Tq-Ca-FLITS. Recently, single FP–based FLIM biosensors were developed for many analytes, including ATP,^36,37^ cAMP,^36^ glucose,^36,38^ lactate,^39,40^ and citrate.^36^ They may have substantial potential for quantifying acute metabolic responses during muscle contraction and systematically evaluating the mechanisms by which exercise, pharmacological, and nutritional interventions regulate muscle metabolism.

In conclusion, we demonstrated the successful introduction of FP–based biosensors for multiple key metabolic parameters, including Ca^2+^, ATP, cAMP, and temperature, into differentiated myotubes, enabling real-time monitoring using FLIM– and intensity-based imaging. These findings confirmed a robust method for multiplexed and real-time analysis of skeletal muscle physiology and metabolism.

## Methods

### Plasmid constructions

Plasmids encoding Tq-Ca-FLIT (# 129628),^12^ MaLionG (# 113906),^13^ ELP-TEMP (# 178434),^15^ Fast-GCaMP6f-RS09 (# 67160)^25^ and Pink Flamindo (# 102356)^14^ were obtained from Addgene (Cambridge, MA, USA). For protein purification, genes encoding the above target biosensors were amplified using PCR with *BamH*I and *Hind*III restriction sites and inserted into a pRSET_A_ expression vector (Thermo Fisher Scientific, Waltham, MA, USA). Additionally, the gene for Tq-Ca-FLIT was amplified with *Nhe*I and *Hind*III and subcloned into a pMax vector (Addgene, # 177825) for mammalian expression.

### Protein expression and purification

Plasmids in the pRSET_A_ vector were transformed into *E. coli* JM109(DE3) competent cells (# P9801, Promega, Madison, WI, USA). The bacteria were cultured in Luria–Bertani medium (200 mL) containing ampicillin (100 µg/mL) at 20 °C with shaking at 120 rpm. Cells were harvested using centrifugation (8000 rpm, 20 min, 4 °C), resuspended in phosphate-buffered saline (PBS; KAC, BNDSBN200) supplemented with a protease inhibitor cocktail (# 11873580001, Roche Diagnostics, Mannheim, Germany), and disrupted using ultrasonication. The lysates were applied to a Ni–NTA agarose column (# 30230, Qiagen, Hilden, Germany), and bound proteins were eluted with a buffer (10 mM Tris–HCl, 150 mM NaCl, pH 8.0) containing imidazole (200 mM; # 099–00013, FUJIFILM Wako Pure Chemical Co., Osaka, Japan). The eluate was subsequently desalted using a PD-10 desalting column (# 17085101, Cytiva, Marlborough, MA, USA) pre-equilibrated with HEPES (20 mM, pH 7.4). The protein solution was concentrated using ultrafiltration with a 50-kDa molecular weight cut-off filter (# UFC803024, Amicon Ultra-4, Sigma-Aldrich, St. Louis, MO, USA).

### Cell culture

C2C12 myoblasts were obtained from American Type Culture Collection (ATCC, Manassas, VA, USA). Cells were seeded in 24-well plates at density of 1.5 × 10^4^ cells/well or 100-mm culture dishes at 1.5 × 10^5^ cells/dish using high-glucose Dulbecco’s Modified Eagle’s Medium (DMEM; # 11965092, Gibco) supplemented with 10% (v/v) foetal bovine serum (# S1400-500, BioWest, Nuaillé, Franc) and 1% (v/v) penicillin-streptomycin (PS). Cells were maintained at 37°C with 5% CO_2_. Upon reaching 80–90% confluence, the culture medium was replaced with differentiation medium (DM) consisting of high-glucose DMEM supplemented with 2% (v/v) calf serum (# 16010159, Gibco), 1% (v/v) PS, and 1% (v/v) non-essential amino acids (# 11140050, Gibco) to induce differentiation into myotubes (designated as day 0). Differentiated myotubes were used for experiments on days 3–14 after induction. The DM was replaced daily.

### Electroporation conditions

Electroporation of the adhered differentiated myotubes was performed using a 4D-Nucleofector^TM^ system (AAF-1003Y, Lonza, Basel, Switzerland) according to the manufacturer’s protocol. The purified Tq-Ca-FLIT proteins were delivered into differentiated myotubes on day 8 for protein delivery, whereas the Tq-Ca-FLIT/pMax plasmid was introduced on day 3 for plasmid delivery. For protein delivery, purified protein solutions were mixed with 4D-Nucleofector^TM^ solution (AD2 4D-Nucleofector® Y Kit, Lonza) to a final volume of 350 μL/well. For plasmid delivery, 32 μL Tq-Ca-FLIT/pMax plasmid (1 μg/μL) was mixed with 350 μL of 4D-Nucleofector^TM^ solution. The electric field strengths were not disclosed by the manufacturers; therefore, preset programs (CA-154, CA-215, FF-166, and FB-191) were used in order of the increasing electric field strength. Simultaneously, 3, 30, and 300 μM Tq-Ca-FLITS proteins were delivered into differentiated myotubes on day 8 to determine the optimal protein concentration. Immediately after electroporation, the cells were washed three times with 1 mL of PBS and cultured in DMEM under 5% CO_2_ at 37 °C overnight. After 24 h or 6–11 days depending on the experiment, the medium was replaced with phenol red-free DMEM/F12 (# 11039-021, Thermo Fisher Scientific) for imaging.

For suspended cells, electroporation was performed using a CUY21EDITII electroporator (BEX Co., Ltd, Tokyo, Japan) following the manufacturer’s protocol. Differentiated myotubes from 100-mm dishes were used for protein and plasmid delivery on day 8 and 3, respectively. Cells were detached using trypsin and resuspended with culture medium. For protein delivery, purified Tq-Ca-FLITS proteins were added to the myotube suspension to a final concentration of 300 µM. For plasmid delivery, 32 μL of the Tq-Ca-FLIT/pMax plasmid (1 μg/μL) was mixed with 350 μL of the myotube suspension. Electroporation was carried out using a 5-ms poration pulse at 75, 125, or 175 V, followed by five 50 ms driving pulses at 50 V. Electroporated myotubes were subsequently seeded onto 35-mm glass-bottomed dishes and cultured under same culturing conditions until imaging.

### Cell imaging with confocal and FLIM

Confocal imaging was performed using an FV1200 confocal microscope (Olympus Co., Tokyo, Japan) equipped with an air objective lens (UPLFLN 10×, NA = 0.3, Olympus). For MaLionG and Fast-GCaMP6f-RS09, excitation was performed at 474 nm and fluorescence emission at 490–540 nm. For Pink Flamindo, excitation was performed at 559 nm and fluorescence emission at 575–675 nm. Images were acquired every 10 s at a resolution of 512 × 512 pixels.

FLIM imaging was conducted using the same FV1200 confocal microscope equipped with a rapid-FLIM^HiRes^ and a MultiHarp 150 Time-Correlated Single Photon Counting unit from PicoQuant. FLIM measurements were performed for Tq-Ca-FLITS and ELP-TEMP. Samples were excited using a 440-nm pulsed laser (PicoQuant GmbH, Berlin, Germany), and fluorescence emission was detected using a 483/40-nm bandpass filter (Semrock, Rochester, NY, USA). The scanning resolution was set at 256 × 256 pixels with a scanning time of 0.429 s/frame. Ten frames were acquired for the fluorescence lifetime analysis. Fluorescence lifetime data were analysed using SymPhoTime 64 software (PicoQuant). All experiments were performed at 23 °C, except for ELP-TEMP experiments, which were conducted at 34 °C. Additionally, an infrared 1470-nm laser (MDL-MD-1470−2W, Changchun New Industries Optoelectronics Tech Co., Ltd., Changchun, China) was installed to directly heat target cells for the ELP–TEMP experiments^41^.

### Cell viability and cytotoxicity

Cell viability and cytotoxicity of electroporated myotubes were evaluated using a Cell Counting Kit-8 (CCK-8; Dojindo Laboratories, Kumamoto, Japan) and Cytotoxicity Lactate Dehydrogenase (LDH) Assay Kit-WST (CK12, Dojindo Laboratories), respectively, according to the manufacturer’s instructions. For the cytotoxicity assay, the culture medium (100 μL) was transferred to a 96-well plate (3860-096, Iwaki, Shizuoka, Japan) and mixed with the LDH working solution (100 μL). The mixtures were incubated in the dark at room temperature for 30 min, followed by the addition of the stop solution (50 μL). The absorbance was then measured at 490 nm using a microplate reader (SpectraMax ABS Plus, Molecular Devices, San Jose, CA, USA). For the viability assay, CCK-8 solution (60 μL) was directly added to each well and incubated for 3 h at 37 °C under 5% CO_2_. The absorbance was subsequently recorded at 450 nm. As a positive control for apoptosis, staurosporine (10 μM; # 569396, Sigma-Aldrich) was added to selected wells.

### Western blotting

Control and electroporated (program: CA-154 and FB-191) myotubes in 24-well plates on day 9 were used for western blotting. To obtain whole protein extracts, cells from 2 wells/group were harvested and lysed in EzRIPA lysis buffer (ATTO Co., Tokyo, Japan) on ice, containing 1% phosphatase inhibitor cocktail (Nacalai Tesque, Kyoto, Japan) and 1% protease inhibitor cocktail (Nacalai Tesque), as previously described.^42^ The whole protein concentration was determined using Protein Assay BCA Kit (Nacalai Tesque). The samples were solubilised in 2× and 1× SDS sample buffers (FUJIFILM Wako Pure Chemical Co.) at 0.5 µg protein/µl and boiled at 95 °C for 5 min.

Whole protein samples were separated using 10% SDS–PAGE, transferred onto polyvinylidene difluoride membranes, and stained with Coomassie Brilliant Blue (CBB) for equal protein loading confimation as previously described.^42^ After the protein transfer, the membranes were blocked in Bullet Blocking One (Nacalai Tesque) for 5 min at room temperature. After several washes with Tris-buffered saline (150 mM NaCl, 25 mM Tris-HCl, pH 7.4) containing 0.1% Tween-20 (TBST), the membranes were incubated overnight at 4 °C with mouse anti-myosin heavy chain (MyHC; 1:2,000, A4. 1025, Developmental Studies Hybridoma Bank, Iowa City, IA, USA) and mouse anti-desmin (1:1,000, DES-DERII-L-CE, Leica Biosystems, Nussloch, Germany) antibodies diluted in TBST containing 5% bovine serum albumin and 0.02% NaN_3_. Then, the membranes were incubated with an anti-mouse IgG horseradish peroxidase-conjugated secondary antibody (1:20,000, NA931V, Cytiva) for 60 min at room temperature. Protein bands were visualised using the ECL Prime reagent (Cytiva) and the signal was captured with MicroChemi (Berthold Technologies, Bad Wildbad, Germany). The band intensities were quantified using ImageJ software (v1.54p; National Institutes of Health, Bethesda, MD, USA). The protein expressions of MyHC and desmin are expressed as a ratio of the values in control group.

### Statistical analysis

All data are presented as mean ± standard deviation. Statistical analyses were performed using Statistical Package for Social Sciences (version 29, IBM Corp., Armonk, NY, USA). Details of the specific statistical tests used are provided in the figure legends. Statistical significance was set at *P* < 0.05.

## Acknowledgements

This work was supported by grant-in-aid from the Ministry of Education, Culture, Sports, Science and Technology, Japan (23KJ1023 to AK; 24H00675 to KM and SA; 25K21726 to SA); the Daiichi-Sankyo “Habataku” Support Program for the Next Generation of Researchers (AK and CQV); Kanazawa University “HOZUMINE” project for promotion of research (TS); JST FOREST Program (JPMJFR201E to SA); and the World Premier International Research Center Initiative (WPI), MEXT. We thank Dr. Djamel Eddine Chafai (WPI-NanoLSI) for supporting electroporation (BEX) machine and Dr. Clemens M. Franz (WPI-NanoLSI) for discussion of muscle physiology.

## Author contributions

A.K. and S.A. conceived and designed the study. A.K., C.Q.V. and N.S. performed the experiments, including electroporation, cellular imaging, and analysed the data. A.K., S.A., C.Q.V. and N.S. performed FLIM analysis. T.S. and K.M. performed protein extraction and western blotting. S.A. and.

C.Q.V. provided biosensors and contributed to methodological development. S.A. and K.M. acquired funding and supervised the project. A.K., C.Q.V. and S.A. wrote the original manuscript. All authors reviewed and approved the final manuscript.

