## Supplemental Data 1 for "Electroporation-mediated delivery of protein biosensors for metabolic imaging in differentiated myotubes"

**Supplementary Information**

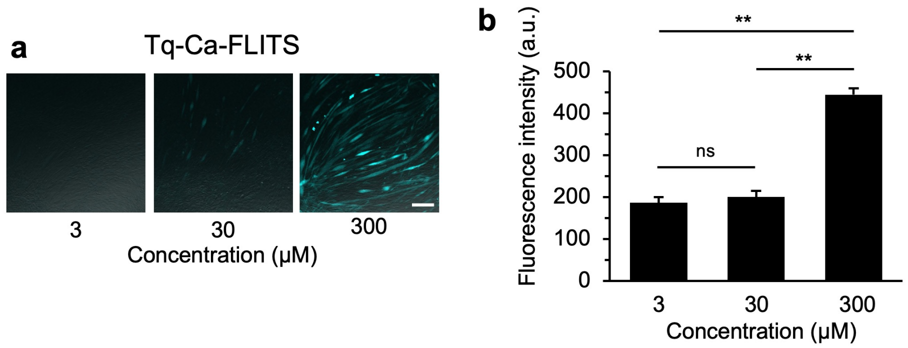


**Supplementary Fig. S1. Effect of protein concentration on delivery efficiency of Tq-Ca-FLITS in differentiated myotubes.** (a) Representative fluorescence images of Tq-Ca-FLITS delivered into differentiated myotubes at increasing protein concentrations (3, 30, and 300 μM). Scale bar, 200 μm. (b) Quantification of fluorescence intensity corresponding to (a). Fluorescence intensity increased with higher protein concentrations, with 300 μM yielding significantly higher signals compared with that for 3 and 30 μM. Data are presented as mean ± SD ($n$ = 12 cells, $N$ = 4). One-way ANOVA followed by Tukey–Kramer post-hoc test was performed. $^{**}P < 0.01$; ns, not significant.

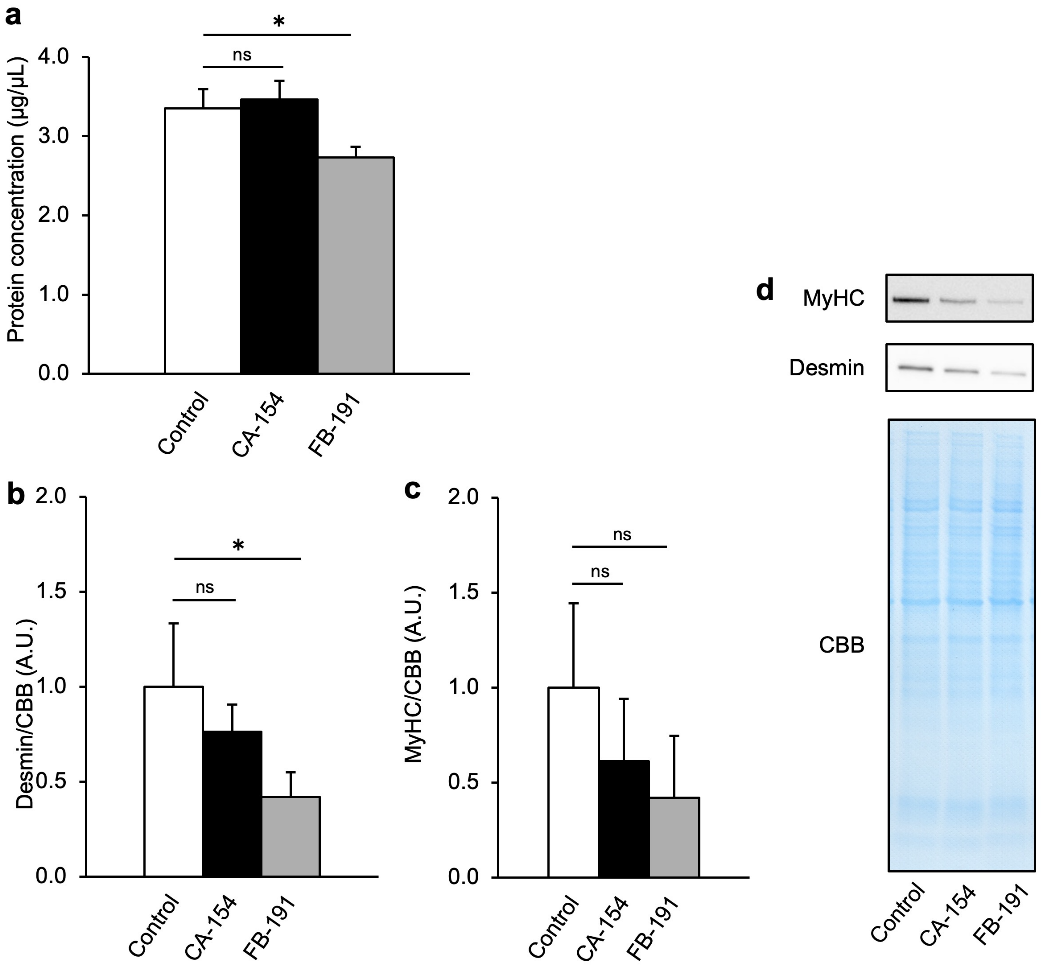


**Supplementary Fig. S2. Assessment of cellular integrity under different electroporation conditions.** (a) Total protein concentration in differentiated myotubes under control (non-electroporated), weak (CA-154), and strong (FB-191) electroporation conditions. (b) Quantification of desmin levels normalised to Coomassie Brilliant Blue (CBB) staining. Desmin levels were reduced under strong electroporation conditions. (c) Quantification of myosin heavy chain (MyHC) levels normalised to CBB staining. No significant differences were observed across conditions. (d) Representative western blot images of MyHC and desmin, along with corresponding CBB–stained gels as loading controls. Data are presented as mean ± SD ($N = 3$). One-way ANOVA followed by Tukey–Kramer post-hoc test was performed. $*P < 0.05$; ns, indicating not significant.

Tq-Ca-FLITS

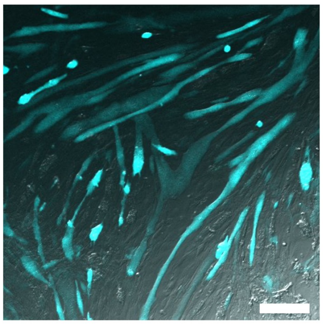

**Supplementary Fig. S3. Rapid intracellular delivery of Tq-Ca-FLITS in differentiated myotubes.** Fluorescence image of Tq-Ca-FLITS delivered into differentiated myotubes within 10–15 min after electroporation. Scale bar, 200 μm.
